# Exact firing time statistics of neurons driven by discrete inhibitory noise

**DOI:** 10.1101/116467

**Authors:** Simona Olmi, David Angulo-Garcia, Alberto Imparato, Alessandro Torcini

## Abstract

Neurons in the intact brain receive a continuous and irregular synaptic bombardment from excitatory and inhibitory presynaptic neurons, which determines the firing activity of the stimulated neuron. In order to investigate the influence of inhibitory stimulation on the firing time statistics, we consider Leaky Integrate-and-Fire neurons subject to inhibitory instantaneous postsynaptic potentials. In particular, we report exact results for the firing rate, the coefficient of variation and the spike train spectrum for various synaptic weight distributions. Our results are not limited to stimulations of infinitesimal amplitude, but they apply as well to finite amplitude post-synaptic potentials, thus being able to capture the effect of rare and large spikes. The developed methods are able to reproduce also the average firing properties of heterogeneous neuronal populations.

## Introduction

Neurons in the neocortex *in vivo* are subject to a continuous synaptic bombardment reflecting the intense network activity^1^. In the so-called *high-input regime*, in which neurons receive hundreds of synaptic inputs during each interspike interval^2^, the firing statistics of model neurons is usually obtained in the context of the *diffusion approximation* (DA)^3,4^.

Within such an approximation the post-synaptic potentials (PSPs) are assumed to have small amplitudes and high arrival rates, therefore the synaptic inputs can be treated as a continuous stochastic process characterized simply by its average and variance, while the shape of the distribution of the amplitudes of the PSPs is irrelevant^5^. However several experimental studies have revealed that rare PSPs of large amplitude can have a fundamental impact in the network activity^6, 7^. Furthermore, the experimentally measured synaptic weight distributions display, both for excitatory and inhibitory PSPs, a long tail towards large amplitudes and a peak at low amplitudes^6–12^. The effect of rare and large excitatory post-synaptic potentials (EPSPs) has been recently examined for generalized leaky integrate-and-fire (LIF) models with generic EPSP distributions^13^ and in balanced sparse networks for conductance based LIF neurons with log-normal EPSP distributions^14^. The presence of few strong synapses induce faster and more reliable responses of the network even for small inputs^13, 14^. Interestingly, in^14^ it has been shown that a single neuron driven by random synaptic inputs log-normally distributed reveals a clear aperiodic stochastic resonance^15, 16^, which is not evident for Gaussian distributed EPSPs.

However, even for the simple case of LIF neurons exact analytic results are still lacking for large PSPs with generic synaptic weight distributions, apart for the case of the exponentially distributed PSPs reported in^17^. In particular, Richardson and Swarbrick have been able to obtain the statistics of interspike interval (ISI) for LIF neurons receiving balanced excitatory and inhibitory Poissonian spike trains with exponentally distributed synaptic weights^17^. Furthermore, results for generic EPSP distributions have been obtained in ^13^ by developing a semi-analytic approach to solve the continuity equation for the membrane potential distribution.

In this paper, we report exact analytic results for the firing time statistics of neurons receiving inhibitory Poissonian spike trains for various synaptic weights distributions. Namely, we estimate the firing time statistics for LIF neurons subject to inhibitory post-synaptic potentials (IPSPs) (with instananeous rise and decay time) characterized by constant amplitude, as well as for uniform and truncated Gaussian IPSP distributions. Furthermore, we apply the developed formalism to sparse inhibitory networks with heterogeneous neuronal properties.

## Models and Methods

### Model and population-based formalism

We will consider the firing statistics of a LIF neuron ^18,19^ subject to a constant external DC current *μ*_0_ and to a synaptic drive *I*(*t*), in this case the dynamical evolution of the membrane potential *v* is given by the following equation

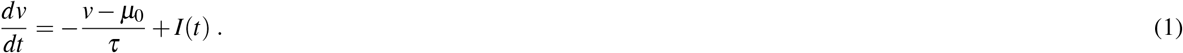

where *τ* = 20 ms is the membrane time constant. The neuron fires whenever the membrane potential reaches the threshold value *v_th_* = 10 mV, afterwards the potential is reset to the value *v_re_* = 5 mV. The synaptic current *I*(*t*) accounts for the linear superposition of the instantaneous excitatory and inhibitory PSPs and it can be written as

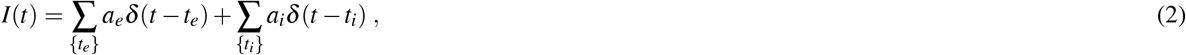

where *a_x_* denote the amplitudes of EPSPs *x* = *e* (*a_e_ >* 0) and IPSPs *x* = *i* (*a_i_ <* 0), while the variables *t_x_* represent their respective arrival times, which are assumed to be Poissonian distributed with rates *R_x_*. Eqs.(1) and ((2)) are equivalent to the Stein’s model^20^ with pulse amplitudes randomly drawn from distributions *A_x_*(*a*). his model, despite its extreme simplicity, has been shown to be able to provide optimal predictions for the average firing rates and spike times of cortical neurons^21,22^.

The aim of this paper is to provide exact analytic expressions for the first two moments of the stationary firing statistics, namely the average firing rate *r*_0_ and the associated coefficient of variation CV, as well as for the spike-train spectrum (STS) *Ĉ*(*ω*)^23^. To obtain such results we follow the approach developed in^17^, in particular within a population-based formalism we introduce the probability density *P*(*v*, *t*) of the membrane potentials together with the associated flux *J*(*v*, *t*). The continuity equation relating these two quantities can be written as:

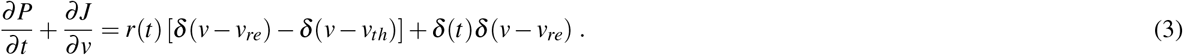

On the r.h.s. of the above equation are reported the sink (source) term for the neuronal population associated to the membrane threshold (reset), with *r*(*t*) being the instantaneous firing rate of the population. The last term on the r.h.s. takes into account the initial distribution of the membrane potentials, which are assumed to be all equal to the reset value at *t* = 0^24^.

The flux *J*(*v*, *t*) can be decomposed in three terms as follows

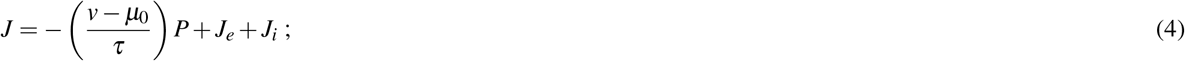

where the first term on the r.h.s is the average drift, while *J_e_* (*J_i_*) represents the excitatory (inhibitory) fluxes originating from the Poissonian synaptic drives. The fluxes can be written as a convolution of the distribution of the membrane potentials with the synaptic amplitude distribution, namely

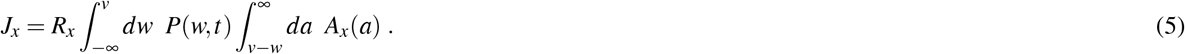

The previous set of equations is complemented by the following boundary conditions:

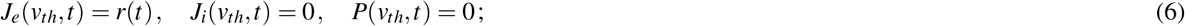

and by the requirement that the membrane potential distribution is properly normalized at any time, i.e

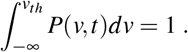

### Analytical method to obtain the exact firing time statistics

The estimation of the firing statistics for a LIF neuron subject to shot noise has proven to be a problem analytically hard to solve^25,26^. The reason is related to the overshoots over the threshold *v_th_* induced by the finite amplitude of the PSPs, which renders difficult the estimation of the membrane potential distribution. However, it is well known that one of the few cases in which the first passage time problem can be solved, is represented by exponentially distributed PSP weights, thanks to the memory-less property associated with exponential distributions^27^. Richardson and Swarbrick^17^ made use of this unique property to derive the exact solution of the firing rate for the case in which both inhibitory and excitatory kick amplitudes are exponentially distributed. The fact that the only boundary relevant for the first passage time is *v_th_*, and considering that no trajectory can cross it from above, implies that inhibitory kicks do not contribute to the overshoot and therefore no restriction over the distribution of their amplitudes should be in principle imposed in order to obtain an analytic solution of the problem.

In the following, using the Laplace transform method, we will derive the analytic expressions for the firing rates, the coefficient of variation and for the spike-train spectrum for various distributions *A_i_*(*a*) of the inhibitory amplitudes. For what concerns the excitatory synaptic input, we will limit our investigation to two analytic solvable cases: namely, to exponentially distributed synaptic weights, where *A_e_* = Θ(*a*) exp(–*a*/*a_e_*)/*a_e_*, and to constant excitatory synaptic drive, encompassed in an external DC current *μ*_0_ *> v_th_*.

Let us first consider the excitatory term for exponentially distributed *a_e_*, in this case the integral equation for the excitatory flux Eq.(5) can be rewritten in a differential form as

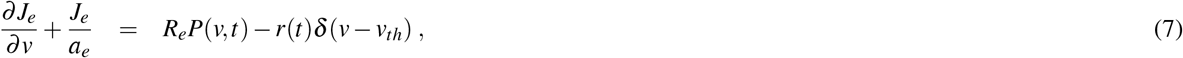

where the last term on the r.h.s. of the above equation accounts for the absorbing boundary condition at threshold. The bidirectional Laplace transform (from now on only Laplace transform) 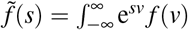 of Eq.(7) can be written as a combination of linear functions in *P̃*(*s*, *t*) and *r*(*t*), namely

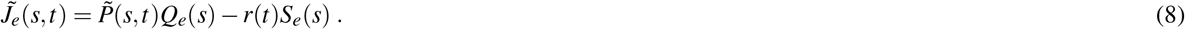

For the particular case of the exponentially distributed EPSP amplitudes

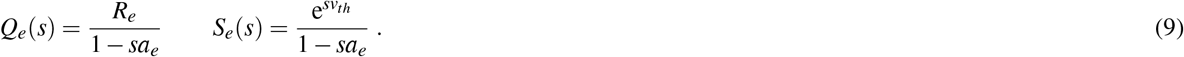

Whenever the excitatory input is simply given by the DC current *μ*_0_, we will assume *J_e_* = *Q_e_* = *S_e_* = 0 and apply the same formulation that we will expose in the following.

For the distribution of inhibitory amplitudes, the only restriction that we will impose is that, one should be able to write the Laplace transform of the inhibitory flux as a linear function of the probability density function, namely *J̃_j_*(*s*, *t*) = *P̃*(*s*, *t*)*Q_j_*(*s*).

### Steady State Firing Rate

Under the above assumptions, we can estimate the Laplace transform of the sub-threshold voltage distribution *Z*_0_ ≡ *Z*_0_(s), which corresponds to the generating function for the sub-threshold voltage moments. Therefore, *Z*_0_ can be estimated as the Laplace transform of *P*(*v*, *t*) when *v_th_* → ∞ implying also that *J* = *r*(*t*) = 0. In particular, by taking the Laplace transform of Eq.(4) together with the assumption that *J* = 0, we obtain

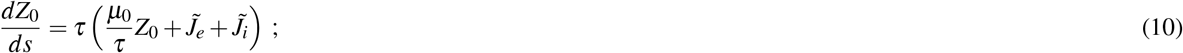

where we set *Z*_0_ = *P̃*.

Since *J̃_e_* and *J̃_j_* are linear in *P̃*, we can rewrite Eq.(10) and solve it as:

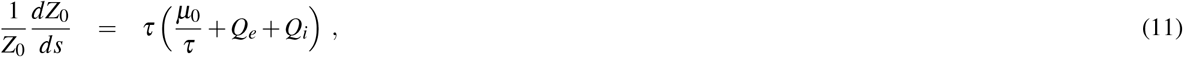

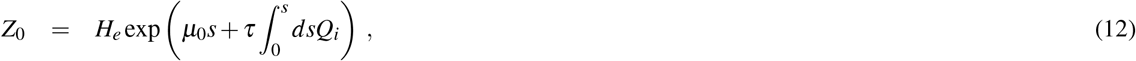

where the excitatory contribution is encompassed in the term

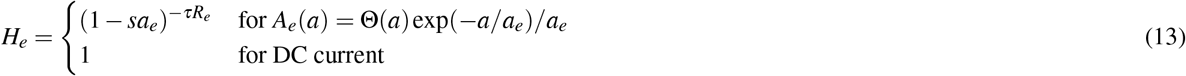

Once we have calculated the generating function, we can solve the stationary case corresponding to *∂P*/*∂t* = 0, performing the Laplace transform of the continuity equation (3), which reads as

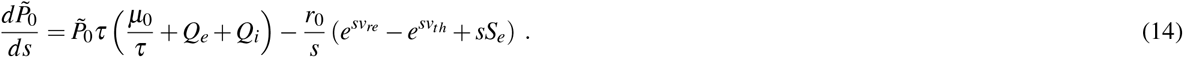

In our notation, the variables with a zero subscript denote stationary quantities. As expected, for *r*_0_ = 0 the function *P̃*_0_ satisfies Eq.(11), therefore we can rewrite the previous equation as

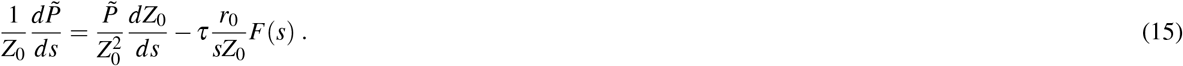

where we have indicated with *F*(*s*) the function multiplying the term *r*_0_/*s* in the r.h.s of Eq.(14). The straightforward solution of Eq.(15) is

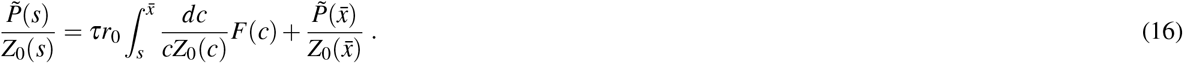

Notice that, although we have derived an expression for *Z*_0_, we have no knowledge of the functional form of *P̃*. Whenever it is possible to identify an integration limit *x̃*, where the term 1/*Z*_0_(*x̄*) vanishes, the following exact analytic expression for the stationary firing rate can be obtained

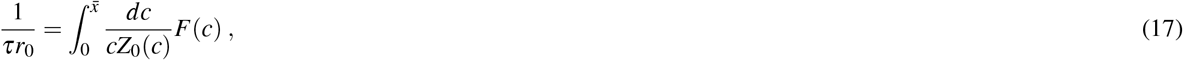

where we have made use of the normalization condition of the probability densities, i.e. *P̃*_0_(0) = Z_0_(0) = 1.

Our analysis is limited to the two previously reported types of excitatory drive, because in these two cases an integration limit *x̄*, where 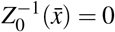, can be easily found to be

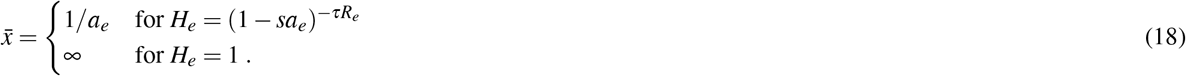

It should be remarked that in presence of both sources of excitatory drive, the integration limit can still be identified whenever *μ*_0_ *< v_th_* and it corresponds to the first one in (18), while we have been unable to solve the case when both, the excitatory drift and the excitatory spike train, can lead the neuron to fire (see conclusions section for a discussion of this point).

### First and Second Moment of the First Passage Time Distribution

Let us now focus on the time dependent evolution of the continuity equation, in this case, the equation can be solved by performing the Fourier transform in time and the Laplace transform in the membrane potential of Eq.(3) and (4). Namely, we obtain

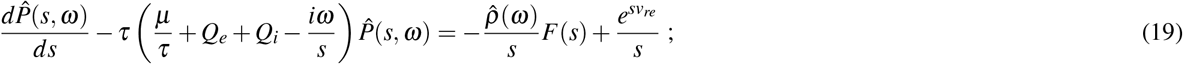

where 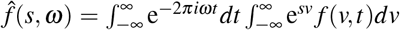 and *ρ̂* (*ω*) is the Fourier transform of the spike-triggered rate. Dividing both sides by *Z*_0_ and integrating the functions over the interval *s* = [0, *x̄*], the l.h.s of Eq.(19) vanishes and an analytic expression for *ρ̂*(*ω*) can be obtained, namely

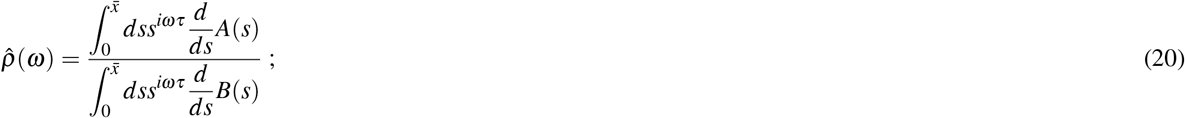

where

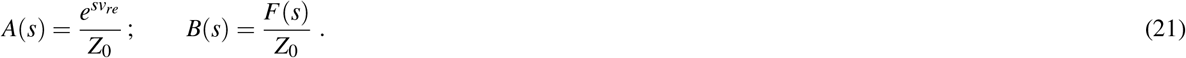

Equation (20) diverges exactly at *ω* ≡ 0, and in that case it should be complemented with the expression *ρ̂*(*ω* = 0) *= r*_0_*πδ*(*ω*). The spike-triggered rate (also called the conditional firing rate) provides all the information on the spike train statistics. For instance, the spike train spectrum *Ĉ*(*ω*), which is the Fourier transform of the auto-correlation function of the spike train, is related to the spike triggered rate via the formula *Ĉ*(*ω*) *= r_0_* [ 1 *+* 2Re(*ρ̂* (*ω*))]^23^.

Moreover, *Ĉ*(*ω*) and *ρ̂* (*ω*) are also related with the Fourier transform of the first-passage time density *q̂*(*ω*) as follows:

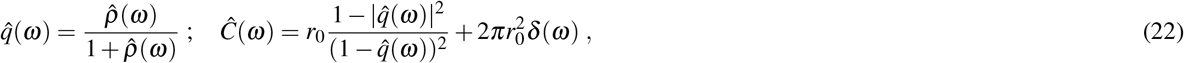

where 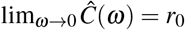 and 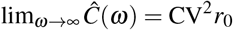^28^.

Therefore the *n* — *th* moment of the first passage time distribution is given by

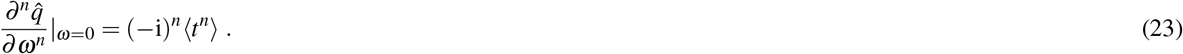

For the estimation of the first two moments it is sufficient to expand to the second order in *ω* the terms entering in Eq.(20), in particular 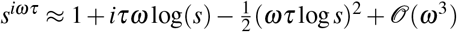. Finally, *ρ̂*(*ω*) can be approximated as a polynomial in ω, that reads as

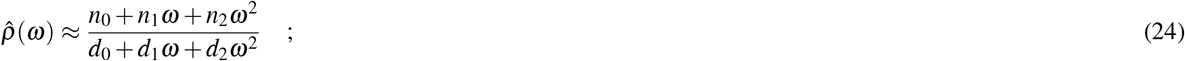

where *n*_0_ = –1, *d*_1_ = *i*/*r*_0_, *d*_0_ = 0 and

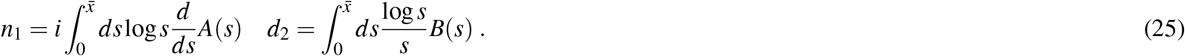

It is worth to notice that, despite the expansion is limited to the second order, the solutions for the first and second moments are exact since terms of higher order disappear when evaluated at *ω* = 0.

From the expression (23) it is easy to verify that the first and second moments of the first passage time distribution are given by

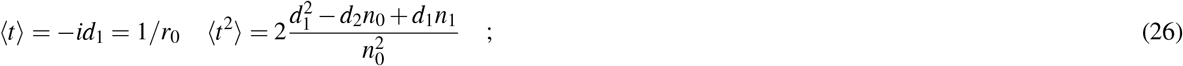

and from these it is straightforward to estimate the coefficient of variation CV of the ISI, namely

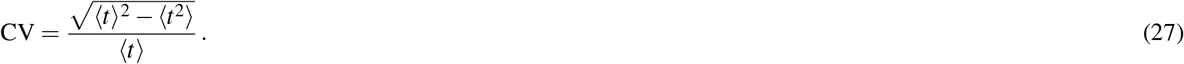

As can be seen by this general solution, the two relevant quantities that define uniquely the stationary firing statistics are the functions *Z*_0_(*s*) and *F*(*s*) defined in the Laplace space, where *F*(*s*) depends only on the considered excitatory drive, while *Z*_0_(*s*) on the whole sub-threshold input.

### Explicit expressions of *Z*_0_ for selected distributions

In this manuscript, we focus on the exact solutions for four relevant types of distributions of the inhibitory synaptic weights *A_i_*(*a*): namely, exponential (ED), *δ* (DD), uniform (UD) and truncated Gaussian distribution (TGD). The exact expressions for *Z*_0_(*s*) for each of these four cases are reported in the following, together with the average and variance of the corresponding distributions *A_i_*(*a*).

#### Exponential Distribution (ED)

In this case the synaptic weight distribution is of the form:

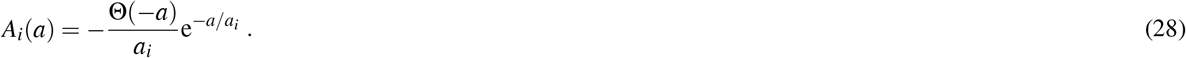

For this distribution, the equations for the inhibitory flux (Eq.(5)) can be rewritten in a differential form analogous to Eq.(7), namely

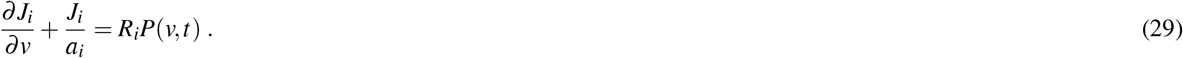

where the term accounting for the absorbing boundary is not present since we are considering the inhibitory shot noise.

The Laplace transform of this equation reads as

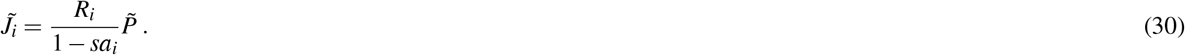

For this choice of distribution, it is possible to obtain an explicit expression for the generating function the sub-threshold voltage moments, namely

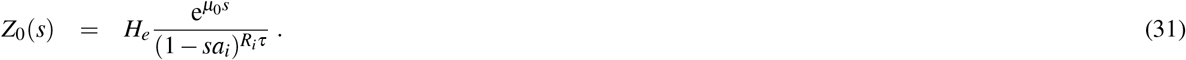

The mean and variance due to the distribution of the inhibitory synaptic weights for the exponential case are 〈*a_i_*〉=*a_i_* and 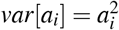 while the absolute value of the skewness is two.

#### δ Distribution (DD)

In the case in which the inhibitory population delivers spikes with a constant amplitude *a_i_*, the distribution becomes simply a *δ*-function:

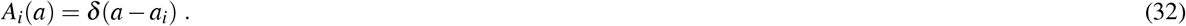

In this case the equation for the inhibitory flux becomes the following

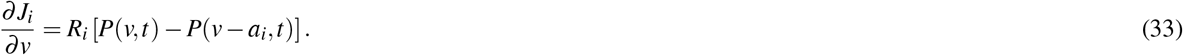

and the associate Laplace transform reads as

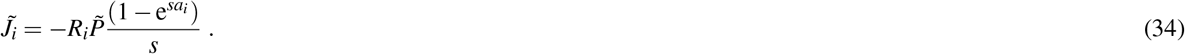

For this simple distribution the generating function *Z*_0_ reads as:

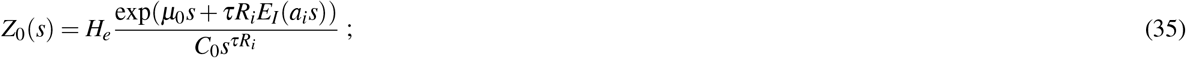

where 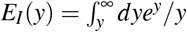 is the exponential integral and *C*_0_ accounts for the normalization condition requiring that *Z*_0_(0) = 1 and its explicit expression is

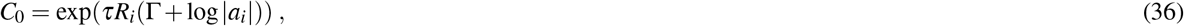

where Γ ≈ 0.577731 is the Euler-Mascheroni constant.

For this distribution the mean and variance of the inhibitory process are simply 〈*a_i_*〉 = *a_i_* and *var*[*a_i_*] = 0, and also the skewness is zero.

#### Uniform Distribution (UD)

For uniformly distributed synaptic weights with support [*l*_1_ *l*_2_], we can express the UD as follows

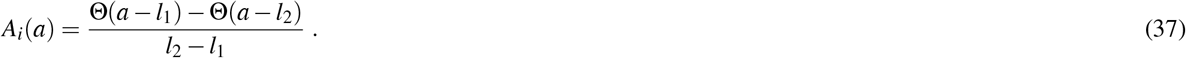

The variation of the flux respect to the potential is given by:

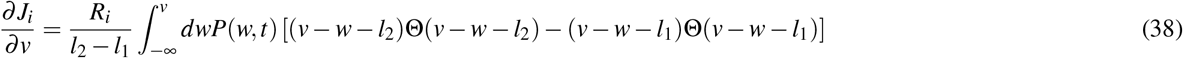

and the corresponding Laplace transform reads as

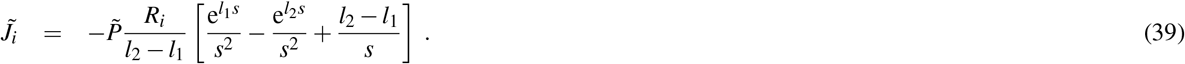

This leads to the generating function together with the normalization constant:

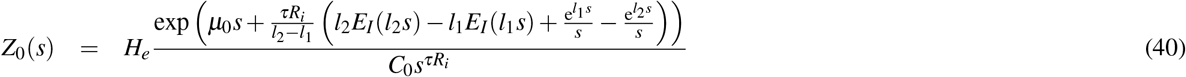

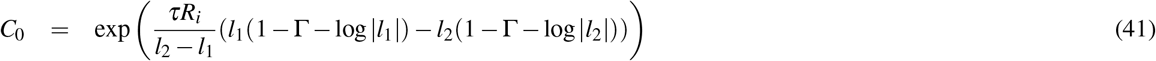

The mean and variance of the inhibitory shot noise are 〈*a_i_*〉 = (*l*_2_ *+l*_1_)/2 and *var*[*a_i_*] = (*l*_2_ *−l*_1_)^2^/12, respectively; while the skewness is zero.

#### Truncated Gaussian Distribution (TGD)

A biologically relevant distribution is the Gaussian distribution^38^, which is peaked at *a_p_* and with a standard deviation equal to *σ_G_*. For notation simplicity we write the Gaussian distribution as

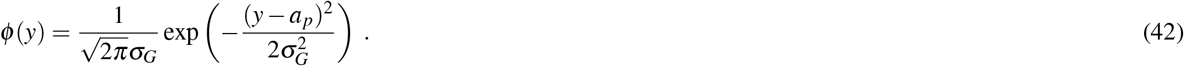

Since we are only interested in the inhibitory kicks, we truncate the original distribution and we impose the support (–∞, 0]. The distribution of the synaptic weights can be written as

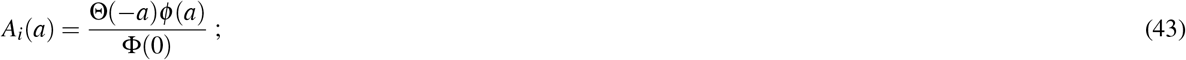

Φ(0) is the cumulative distribution of the normal distribution evaluated at the upper limit of the support according to the equation

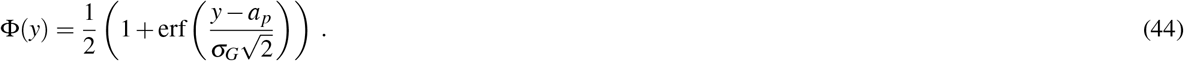

The flux of inhibitory probability takes the form

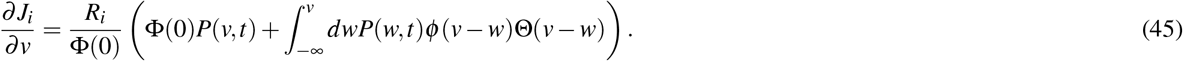

The corresponding Laplace transform is:

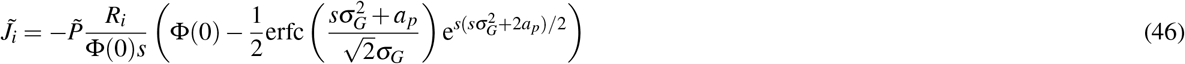

The complicated expression appearing in Eq.(46) does not allow us to obtain the explicit expression for *Z*_0_. However it is easy to integrate numerically Eq.(12) once we know 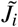. When dealing with a width of the Gaussian distribution *σ_G_ >* 1, the exponential term in Eq.(46) can grow very rapidly while the complementary error function tends to 0, generating numerical problems due to machine precision, specially in the evaluation of the terms *s* > 1. In such cases one can use the first order expansion of the complementary error function: 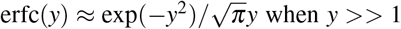 and Eq.(46) can be simplified in such cases to:

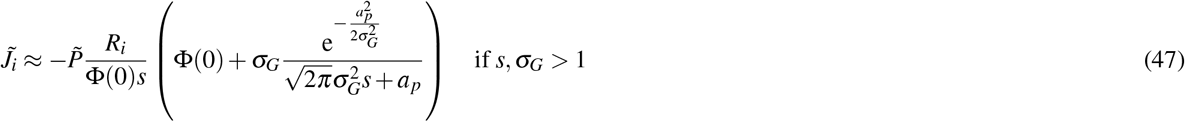

For the TGD, the mean value and variance of the membrane potentials associated to the inhibitory shot noise take the following form

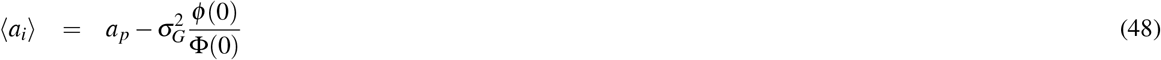

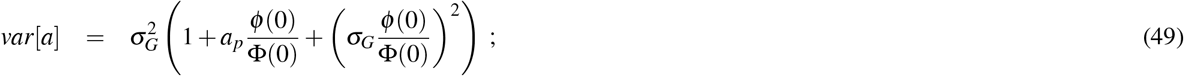

The value of the negative skewness grows with *σ_G_*, namely it passes from ≃ –0.22 for *σ_G_* = |*a_i_*/2| to ≃ −0.92 for *σ_G_* = |5*a_i_*| and it tends to the value 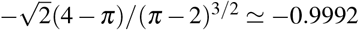 for *σ_G_* → ∞.

### Effective Input and Synaptic Noise Intensity

In order to perform meaningful comparisons between the neuronal response for different synaptic weights distributions, we will consider the responses obtained for the same *effective* average input *μ_T_* and noise intensity *σ*.

For a neuron receiving an inhibitory Poissonian spike train at a rate *R_i_* and with an average synaptic weight 〈*a_i_*〉 the effective average input is

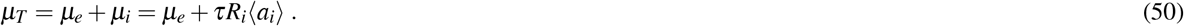

where *a_i_* < 0 and *μ_e_* is the average excitatory input. When only the drift term is present *μ_e_* = *μ*_0_, while for a neuron receiving also an excitatory Possonian spike train of rate *R_e_* and with average synaptic weight 〈*a_e_*〉, it becomes *μ_e_* = *μ_0_* + *τ R_e_* 〈*a_e_*〉. On the one hand, for an effective sub-threshold input *μ_T_* < *v_th_* the dynamics of the LIF neuron is characterized by two timescales: the relaxation time from the reset value *v_re_* to the resting value *μ_T_* and the activation time associated to the escape process from the resting state to the threshold induced by the fluctuations in the input^28^. On the other hand, for a supra-threshold LIF neuron for which *μ_T_* > *v_th_*, in absence of a refractory state, the only characteristic time is the tonic firing rate

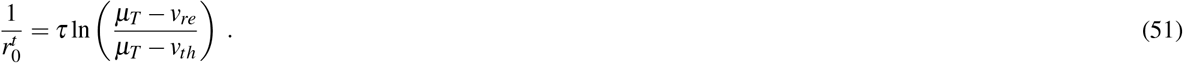

Moreover, in the set-up that we are studying, the neuron is in general subjected to two sources of randomness. A first source due to the variability of the arrival times of the Poisson process, and a second source due to the distribution of the amplitudes of the synaptic weights. Therefore, the total noise intensity *σ*^2^ associated to these two uncorrelated processes is given by the sum of the variances of each process, namely

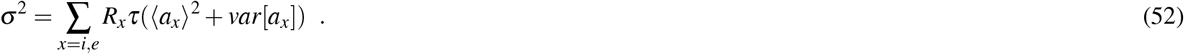

From Eqs.(50) and (52) one can see that the values of *μ_T_* and *σ* can be independently tuned with an appropriate selection of the parameter *μ_e_*, and *R_i_* for any fixed average inhibitory amplitude. Whenever the excitatory drive is given simply by a DC term *μ*_0_, for any choice of the couple of values {*μ_T_*, *σ*} one has an unique solution, which however does not always correspond to a firing activity. For instance, at small values of *σ*^2^ one can find values of *μ*_0_ < *v_th_*, meaning that the neuron will never fire. This implies the existence of a firing onset threshold for this class of systems which depends on the shape of the distribution. Conversely, when exponentially distributed excitatory kicks are present, two parameters (amplitude and the rate of arrival of the excitatory kicks) define *μ_e_* and therefore a whole family of solutions exist for any fixed *a_i_*. Therefore in this case there is not a finite firing onset threshold, allowing for arbitrarily small rate even for small noise intensities.

Throughout this paper we will compare the results obtained within the shot noise framework against the widely used diffusion approximation^3^. Within such approximation the particular shape of the synaptic weight distributions are irrelevant and indeed the only relevant quantities defining the stationary firing rate and the CV are *μ_T_* and *σ*^2^ through the formulas

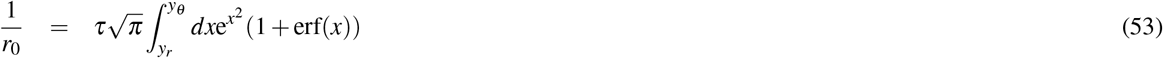

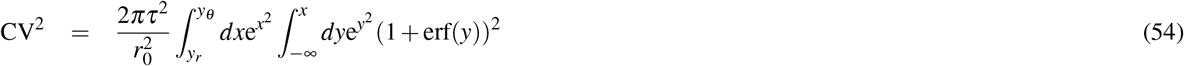

where *y_θ_* = (*v_th_* – *μ_T_*)/*σ* and *y_r_* = (*v_re_* – *μ_T_*)/*σ*.

By following^17^, Eqs.(53) and (54) can be obtained from the shot noise formulation, namely from Eq.(26) and Eq.(22) by considering the expansion of *Z*_0_ limited to the first two voltage moments:

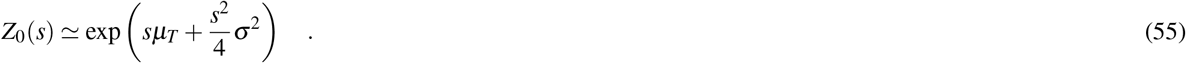

## Results

In this Section we will apply the developed formalism to estimate the response of a single LIF neuron as well as the firing characteristics of a sparse inhibitory neural network for different synaptic weight distributions. In particular, to verify the limits of applicability of the approach we will compare the theoretical estimations with numerical data and with the DA.

Usually, the firing time statistics of a neuron subject to a noisy uncorrelated input is theoretically estimated within the so-called DA^3,4,29^. This approximation is only valid, however, when the PSP amplitudes are small compared with the reset-threshold voltage distance and the arrival frequencies are sufficiently high. Outside of such limits the DA fails to reproduce the numerical data and in particular it is unable to capture the differences due to different synaptic weight distributions^13,17^. Furthermore, the DA has been employed to reproduce network activity of recurrent networks, in such a case one should assume that the spike trains impinging on the neuron are temporally uncorrelated, a condition usually fulfilled in sparse networks^29,30^. However this should be considered as a first approximation, indeed correlations are present even in sparse balanced networks and they can be captured by driving a single neuron with a colored Gaussian noise self-consistently generated^31^.

### Influence of the IPSP distributions on the firing statistics

As a first aspect, let us consider the dynamics of a single LIF neuron subject to an excitatory DC current *μ_e_* = *μ*_0_ > *v_th_* plus the inhibitory contribution given by a Poissonian train of IPSPS with constant amplitude *a*_*i*_. In particular, we examine the neuronal response by increasing the noise intensity *σ*^2^ at a constant effective input current, namely *μ_T_* = 9 mV, corresponding to a sub-threshold case. In particular, the noise intensity is varied by adjusting the rate of the inhibitory train *R_i_*. The results, reported in Fig. 1 (a), show that the neuron fires only for sufficiently large noise and the firing rate *r*_0_ increases with *σ*^2^ as expected. Furthermore, as shown in Fig. 1 (b) the coefficient of variation exhibits a clear minimum at an intermediate noise amplitude 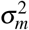, an effect known as coherence resonance (CR) and widely studied in the context of excitable systems^32^. The emergence of CR is related to the presence of at least two competing time scales which depend differently on the noise amplitude, for the sub-threshold LIF neuron these two time scales are the relaxation and escape times^28^.

**Figure 1.**
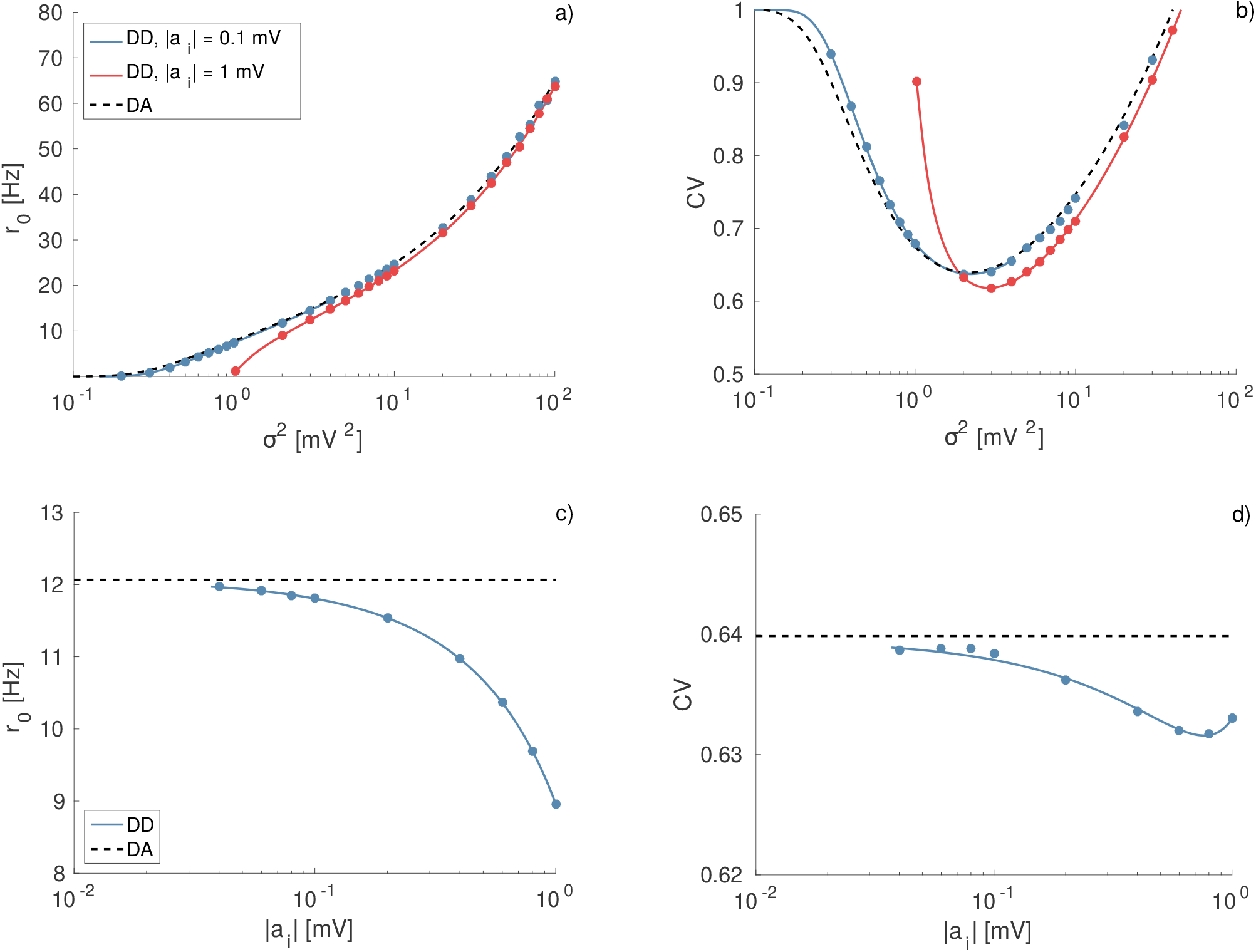
Firing time statistics for different IPSP amplitudes with *δ*-distributions (DD). Average firing rate *r*_0_ (a) and coefficient of variation CV (b) as a function of noise intensity *σ*^2^. The red curves/symbols correspond to |*a_i_*| = 1 mV and the blue ones to |*a_i_*| = 0.1 mV. Firing rate *r*_0_ (c) and CV (d) as a function of the synaptic amplitude |*a_i_*| for constant noise intensity, namely *σ*^2^ = 2 mV^2^. Black dashed curves refer to the diffusion approximation (DA) (Eqs.(53) and (54)); solid curves to the theoretical results for shot noise with constant amplitude *a_i_* (Eqs.(17) and (27)) and symbols to the corresponding numerical simulations. Simulations were performed by exactly integrating Eq.(1) with an event driven scheme (see^34^ for details) and the statistics were estimated over ≈ 10^7^ spikes. For all the panels the effective input is *μ_T_* = 9.0 mV and the excitatory drive is simply a DC term; other parameter values are *v_re_* = 5 mV, *v_th_* = 10 mV, *τ* = 20 ms.

For sufficiently small IPSP amplitudes the agreement between the DA, the numerical results and the analytic expression reported in Eq.(17) is almost perfect as evident from Fig. 1 (a-b), where the blue symbols/curve refer to |*a_i_*| = 0.1 mV. However, by increasing the IPSP amplitude the agreement between numerical results and the estimation given by the shot-noise result (17) remains very good, while the DA is unable to capture the effect of large IPSP. In particular, for |*a_i_*| = 1 mV the DA fails in reproducing the onset of the firing activity as well as the position and height of the minimum of the CV, and in general the firing statistics for low noise amplitudes (see red curves/symbols in Fig. 1 (a-b)). Unitary IPSPs of amplitude ≃ 1 mV have been measured experimentally in the hippocampus of Guinea-pig *in vitro*^8^.

As shown in Fig. 1 (a-b), the increase of the IPSP amplitude has a noticeable effect on the neuronal response, even if the effective input *μ_T_* and the noise intensity are the same as in the case |*a_i_*| = 0.1 mV. Increasing IPSP amplitudes leads to a decrease in the firing activity and it induces an increase of the maximal coherence observable at 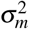, which now occurs at larger noise amplitude with respect to the case |*a_i_* | = 0.1 mV. This can be explained by the fact that the relaxation times to the equilibrium value *μ_T_* are longer for larger IPSP and this induces a deeper minimum in the CV as reported in^33^. Furthermore the irregularity in the emitted spikes, as measured by the CV, increases for 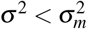 and becomes more regular at larger noise intensities.

The effect of the IPSP amplitude can be better appreciated by performing a different test, namely by maintaining constant both the noise intensity and *μ_T_* while increasing |*a_i_*|. These two quantities can be independently tuned with a suitable selection of *R_i_* and *μ*_0_ in Eqs.(50) and (52). The results of this analysis are reported in Fig. 1 (c-d), the firing rate exhibits a dramatic decrease for increasing |*a_i_*|, as expected due to the increase of average inhibitory current *μ_i_*. This effect is well reproduced by Eq.(17), but it is absolutely not captured by the DA as shown in Fig. 1 (c). The increase of the IPSP amplitude leads to a small variation of the CV revealing a minimum at an intermediate value |*a_i_*| ≃ 0.9 mV (see Fig. 1 (d)). Nevertheless, the DA provides a constant value for the CV in the whole examined range. The origin of the minimum can be understood observing Fig. 1 (b): by increasing |*a_i_*| the overall minimum of the CV curve shifts towards larger noise amplitudes, and at the same time the CV values decrease (increase) for 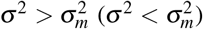). In Fig. 1 (d), we consider a noise intensity *σ*^2^ = 2 mV^2^ for all the simulations. For small (large) |*a_i_*| the maximal coherence is observable at 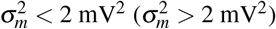), thus the CV value at *σ*^2^ = 2 mV^2^ decreases (increases) with |*a_i_*|. The minimum in Fig. 1 (d) occurs exactly when the CV displays its absolute minimum at 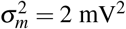.

For the moment we have considered only the case of constant IPSP amplitudes, now we will examine the influence of different distributions *A_i_* on the response of the single LIF neuron. In general, we observe that the shape of the IPSP distribution can noticeably influence the firing rate and the CV. In order to verify this observation, we consider the firing statistics of a neuron subject to the same average input and noise intensity obtained by considering inhibitory spike trains with IPSP distributions of different shapes, but with the same average amplitude 〈|*a_i_*|〉. In particular, more asymmetric distributions, characterized by higher skewness and presenting longer tails towards larger IPSP amplitudes, induce lower firing rates, as shown in Fig. 2 (a) and (c) for supra-threshold and sub-threshold neurons, respectively. In the sub-threshold case this implies that passing from *δ*-distributions, to uniform, to truncated Gaussian and to exponential ones, the firing onset will occur for larger and larger *σ*^2^. For sub-threshold neurons the finite IPSP amplitude enhance the coherence resonance effect with respect to infinitesimal IPSP (corresponding to the DA), while the long inhibitory tails induce a shift of the minimum in the CV towards larger noise amplitudes (see Fig. 2 (b)). For supra-threshold neurons the increased asymmetry in the distributions simply induces more regular firing, as shown in Fig. 2 (d). It is quite peculiar that the TGD and the UD give almost identical results, despite the fact that the TGD is more asymmetric and characterized by a larger standard deviation, namely 0.61 mV for the TGD and 0.29 mV for the UD. Differences are instead seen with respect to the ED which reveals extremely long tails and a standard deviation of 1 mV and to the DD where there is no variability in the IPSP amplitude.

**Figure 2.**
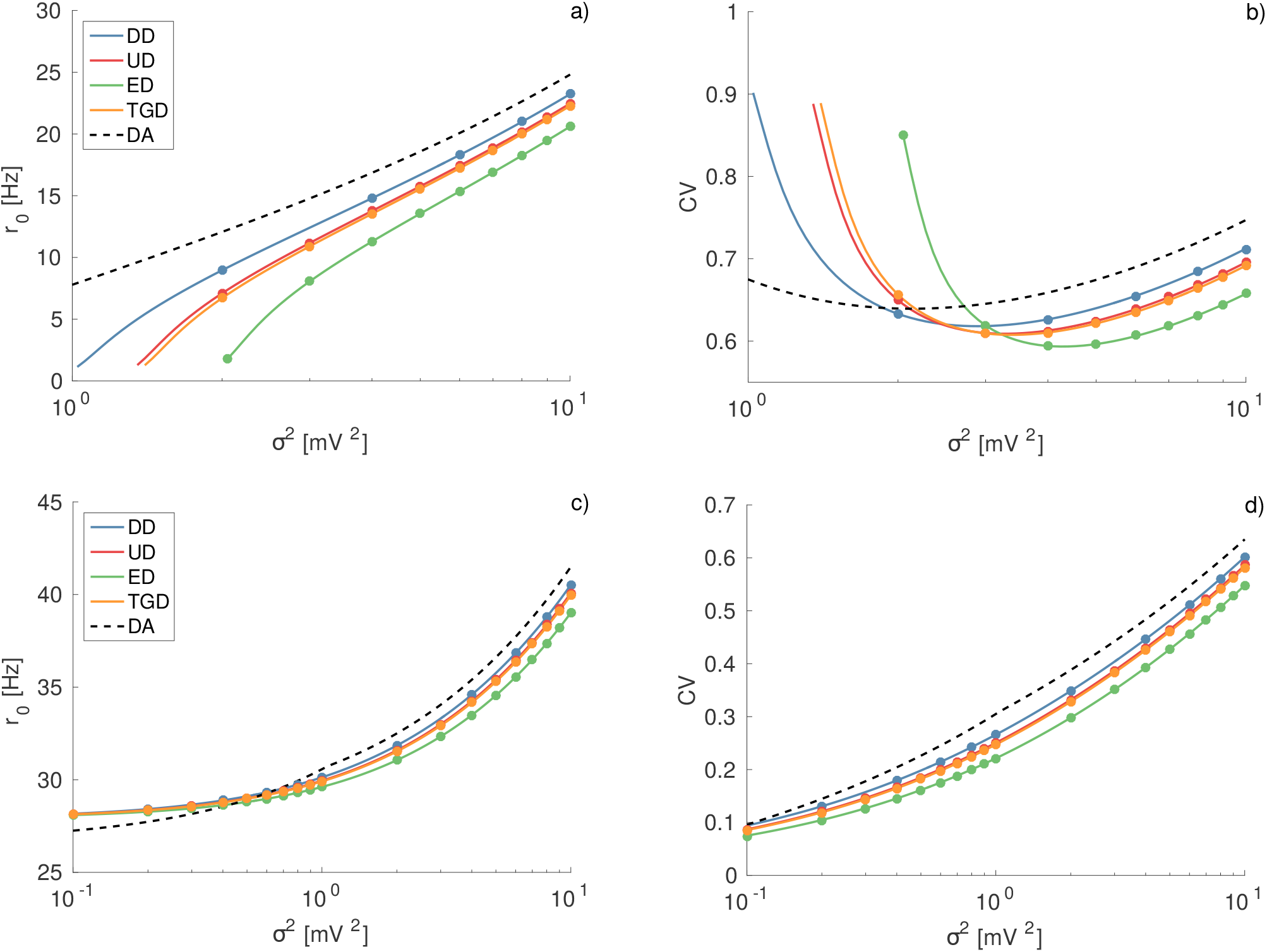
Firing time statistics for different IPSP distributions. Firing rate *r*_0_ and CV as a function of noise intensity for sub-threshold (*μ_T_* = 9 mV) (a,b) and supra-threshold neurons (*μ_T_* = 11 mV) (c,d). In all the panels the symbols correspond to numerical simulations, the dashed black lines to the DA, calculated from Eqs.(53) and (54), and the solid lines to the theoretical results reported in Eq.(17) and Eq.(27). The average IPSP amplitude is set to 〈|*a_i_*|〉 = 1 mV for all the distributions. For the TGD: the peak position |*a_p_*| and the width *σ*_*G*_ of the distribution are equal (namely, ≈ 0.7766 mV). For the UD: *l_1_* = 2*a_i_* and *l*_2_ = 0. Other parameters and simulation procedures as in Fig. 1.

A much more detailed characterization of the firing time statistics, beyond the first two moments that we have considered so far, can be achieved by evaluating the spike train spectrum *Ĉ*(*ω*) which is directly related with the first-passage time density (22). The comparison of the theoretical estimations with the numerical findings is reported in Fig. 3, showing a very good agreement for all the reported cases. We report each spectra normalized by the corresponding average firing rate *r*_0_, in order to emphasize the changes produced by the shape of the IPSP distribution, rather than the changes due to the different values of the firing rates. From Fig. 3 (a), we observe that for *δ*-distributed synaptic weights, the increase in the kick amplitude *a_i_* induces a higher peak in the spectrum at a lower frequency. Therefore, for increasing *a_i_* not only the rate decreases, as previously reported, but also the peak of the ISI distribution shifts from 41 msec for |*a_i_*| = 0.1 mV to 54 msec for |*a_i_*| = 1 mV, thus suggesting that the entire dynamics slows down for increasing IPSP amplitudes.

**Figure 3.**
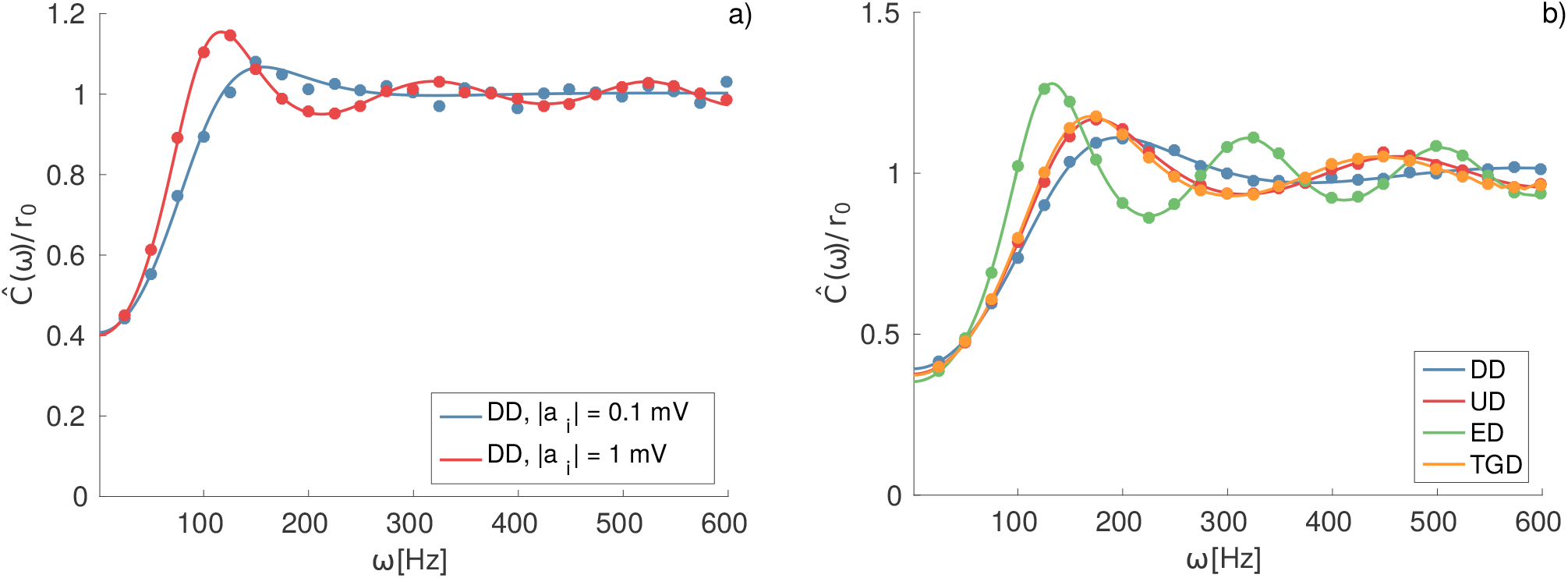
Spike Train Spectra. a) Normalized spike train spectra *Ĉ*(*ω*)/*r*_0_ for the same cases shown in Fig. 1 (a,b) for an effective input *μ*_*T*_ = 9 mV and noise intensity *σ*^2^ = 2 mV^2^; b) Normalized spike train spectra for the distributions considered in Fig. 2 (a,b) for 〈|*a_i_*|〉 = 1, *μ_T_* = 9mV and *σ*^2^ = 4 mV^2^. In both panels, the symbols correspond to the simulation results, while the continuous line to the theoretical estimations using Eq.(22). Simulated spectra were obtained by calculating the squared modulus of the Fourier transform of the spike train using a time trace of 64 s with 1 ms binning, and averaging over 10,000 realizations.

Furthermore, as shown in Fig. 3 (b) the shape of the distributions of the IPSPs has also an influence on the spectra. In particular, the increase in the asymmetry of the distributions induces peaks at lower and lower frequencies, while their height increases. The neuron activity is slowed down for longer and longer tails in the IPSP distributions.

These conclusions are supported by the data reported in Fig. 4 (a). In the figure are reported the average firing rate *r*_0_ for TGDs with different standard deviation *σ_G_* and fixed position of the maximum at *a_p_* = –1 mV. For increasing *σ_G_* the neuronal activity decreases for corresponding noise amplitudes. Since the value of the skewness increases with *σ_G_*, more asymmetric IPSP distributions induce a lower neuronal firing rate. Furthermore, a larger asymmetry is also responsible for a shift of the position of the minimum of CV, associated to the CR phenomenon, towards larger *σ*^2^ and for a more regular firing activity observable at 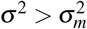, as shown in Fig. 4 (b).

**Figure 4.**
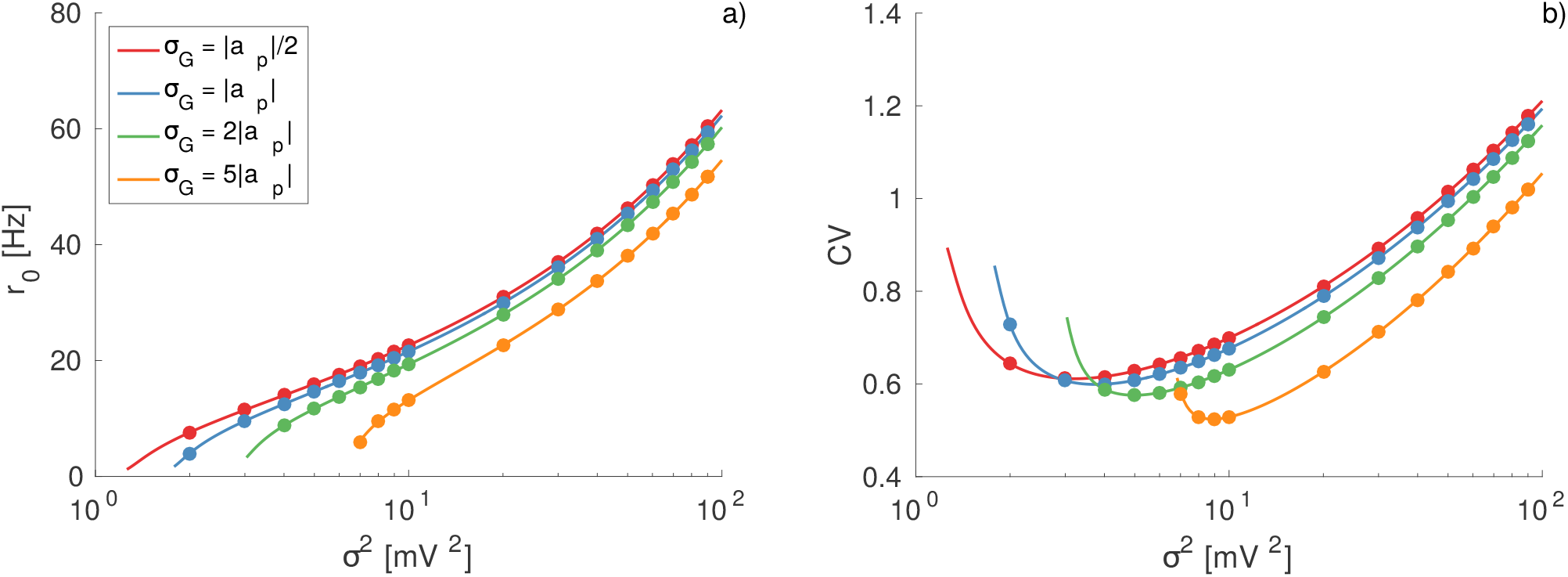
Effect of the distribution asymmetry on the firing time statistics. a) Average firing rate *r*_0_ as a function of the noise intensity for different values of the standard deviation of the Gaussian distribution *σ_G_* as indicated in the legend. b) Coefficient of variation CV for the same cases depicted in a). The reported data refer to TGDs with the peak located at |*a_p_*| = 1 mV and to an effective input *μ_T_* = 9 mV. Symbols correspond to numerical simulations, and the solid lines to the theoretical results reported in Eq.(17) and Eq.(27). Other parameters and simulation procedures as in Fig. 1

So far we considered as excitatory input a constant DC term, however the analytic approach here presented can be applied also for exponentially distributed excitatory amplitudes, namely *A_e_* = exp(*–a*/*a_e_*)/*a_e_*. We have reported the results for this case for different inhibitory distributions in Fig. 5, the analytic estimations reproduce very well the numerical results both for the firing rate and the CV in an ample range of noise intensities. However, for small values of the noise intensity, the analytic results slightly deviates from the numerical values due to the fact that in the limit of small rates the numerical evaluation of the integrals entering in the expresssion of the generating function *Z*_0_ have problems of convergece. The effect due to the different IPSP distributions is analogous to the one observed with a constant DC excitatory term. The DA, conversely, is unable to capture the precise values of these two quantities. Once more, for the reported case in Fig. 5 where *μ_T_* = 9 mV, the CR effect is present. It is important to remark that, in the specific case of exponentially distributed amplitudes of the EPSP, the approach discussed in this article fails when an additional supra-threshold DC current is applied, namely for *μ*_0_ > *v_th_*. A brief discussion in this regard will be provided in the concluding section.

**Figure 5.**
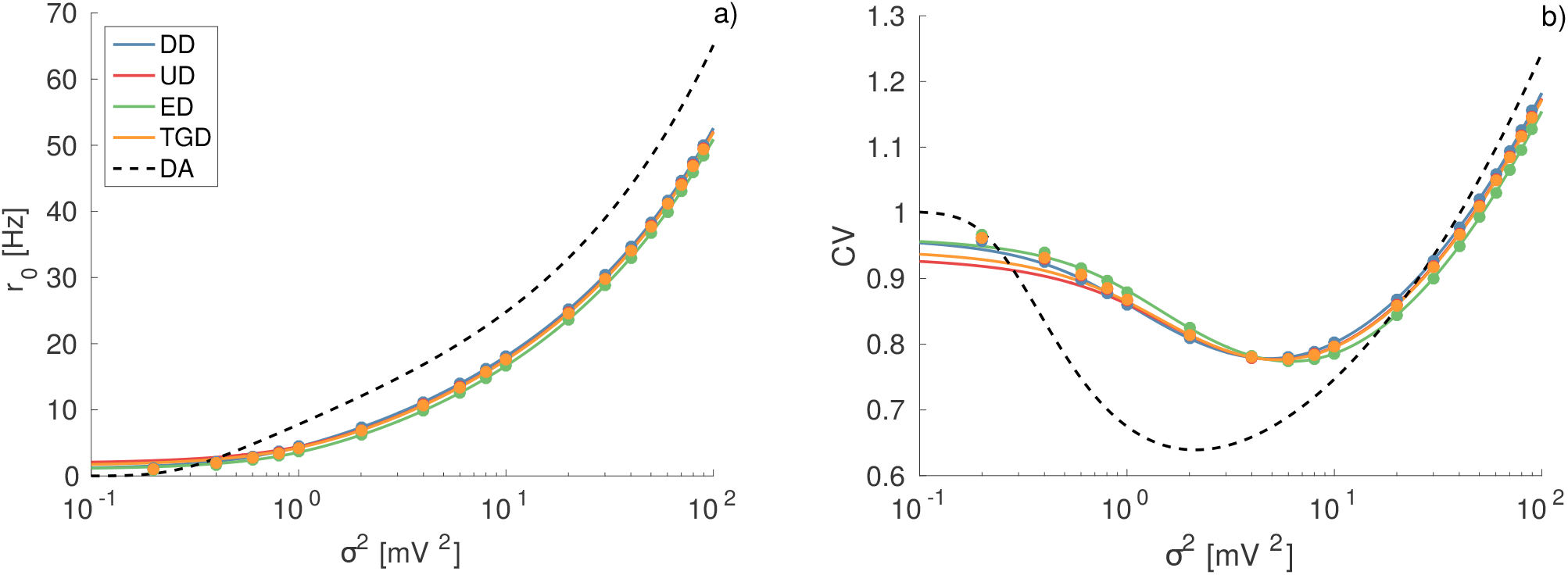
Firing time statistics for exponentially distributed EPSPs. Firing rate *r*_0_ (a) and CV (b) as a function of noise intensity for excitatory synaptic weights distributed exponentially with average amplitude 〈*a_e_*〉 = 〈|*a_i_*|〉 = 1 mV. In this figure excitatory and inhibitory spike trains are balanced, i.e *R_e_* = *R_i_* while the effective input is sub-threshold, namely *μ_T_* = *μ*_0_ = 9 mV. The symbols and lines have the same definition as in Fig. 2, as well as all the other parameters for the IPSP distributions and the numerical simulations.

### Heterogeneous Sparse Networks

The previous theoretical analysis of single neuron response to an external Poissonian input can find application also in the analysis of the dynamics of recurrent LIF heterogeneous networks with random sparse connectivity. For sparse networks, the spike-trains impinging a certain neuron can be assumed to be uncorrelated and Poissonian^29,35^. Furthermore, similarly to what done in ^29^, we can assume that the spike-trains in the networks can be self-consistently described as Poissonian processes with firing rates *r*_0_(*μ*(*j*)) related to the neuronal excitability *μ*(*j*) of each single neuron.

For the sake of simplicity, we will consider a network of inhibitory neurons, where each neuron is characterized by a different level of excitability {*μ*(*j*)} encompassing any excitatory external drive as well as the specific characteristic of the considered neuron. Therefore, the dynamics of the *j*-th neuron in the network can be written as 
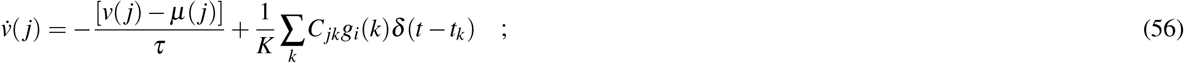

where *K* << *N* is the number of synaptic neighbors, *C_jk_* is the connectivity matrix with entries 1 (0) if the *k*-th neuron is connected (not connected) to neuron *j*. The amplitudes of the IPSP *a_i_*(*k*) = *g_i_*(*k*)/*K* < 0 associated to the firing of the k-th neuron is assumed to be randomly distributed following some of the distributions previously introduced.

For the population dynamics we will give an estimate of the average firing rate via a self-consistent approach by assuming that in average each neuron receive a single Poissonian spike train with a rate *R_i_* = *r̄*_0_*K* and where each IPSP has a random amplitude *a_i_* taken from a distribution *A_i_*(*a*). An estimation of the average firing rate can be obtained by solving the following implicit equation

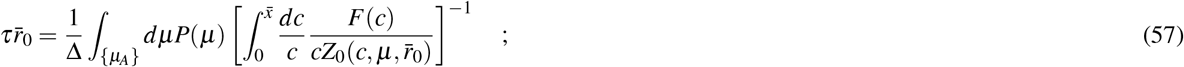

which is an extension to an heterogeneous network of Eq.(17) and where, as we have previously discussed, *F*(*c*) depends on the excitatory input and *Z*_0_ on the chosen IPSP distribution. It is important to stress that usually in inhibitory networks not all neurons are firing, but just a certain fraction *n*\* will be active^34^, therefore the integral reported in (57) is limited to these neurons, which are the one with higher excitability *μ*(*j*) ∈ {*μ_Α_*}. Furthermore Δ = *∫*_{*μA*}_ *dμP*(*μ*) is the support of the active neurons. Once the firing rates have been obtained self consistently we can derive the coefficient of variation of each single neuron according to Eqs.(26) and (27) and then to perform the population average as follows

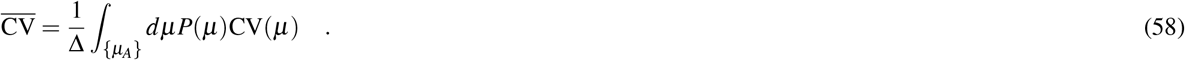

In^34^ a theoretical approach to obtain self-consistently *n*\*, 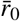 and the average coefficient of variation 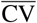 has been developed for constant IPSP, i.e. for *δ*-distribution. Here we extend such approach to more generic IPSP distributions, however we will limit to obtain the analytic estimations of 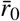 and 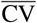. The values of *n*\*, entering in the expressions for the average population rate and coefficient of variation, will be considered as parameter values obtained directly from the simulations. We have verified that the comparison with the numerical findings is very good for all the four considered IPSP distributions and for the whole considered range of average synaptic inputs 〈*a_i_*〉. For illustration purposes we present in Fig. 6 (a) and (b) the average firing rate 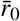 and 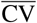, for the two distributions that have consistently presented larger differences between them, namely DD and ED, and for the DA. While it is evident that the DA overestimates 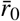 and 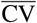 already for 〈|*a_i_*|〉 = 0.5 mV, we observe that in the case of heterogeneous networks the differences among the various IPSP distributions are quite limited at the level of the average firing rate 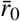 and 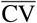. In order to investigate more in details the influence of different IPSP distributions on the neuronal response, we have numerically estimated the probability distribution functions *P*(*r*_0_) of the single neuron firing rate *r*_0_ for DD and ED distributions. As shown in Fig. 6 (c) and (d), the *P*(*r*_0_) obtained for the DD display a higher peak at low firing rate *r*_0_ with respect to the ED, while for higher firing rates they essentially coincide. This difference should be ascribed to the fact that sub-threshold neurons subject to IPSP trains with ED have a firing onset at definitely larger noise amplitudes than the ones subject to IPSPs with DD, while for supra-threshold neurons the differences are definitely less evident, as previously reported in Fig. 2 (a) and (c). This explains also why the exam of the average firing rate does not reveal large difference between the two distributions, since the neurons with low firing rate contribute negligibly to the average activity. The influence of synaptic weight distributions on neuronal population dynamics has been usually examined in the context of homogeneous neuronal populations^13,17^, where the single neuron response represents a good mean field approximation of the network dynamics. However, from the present analysis it emerges that the heterogeneity in the single neuron excitability can render extremely difficult to distinguish among stimulations with different IPSP distributions.

**Figure 6.**
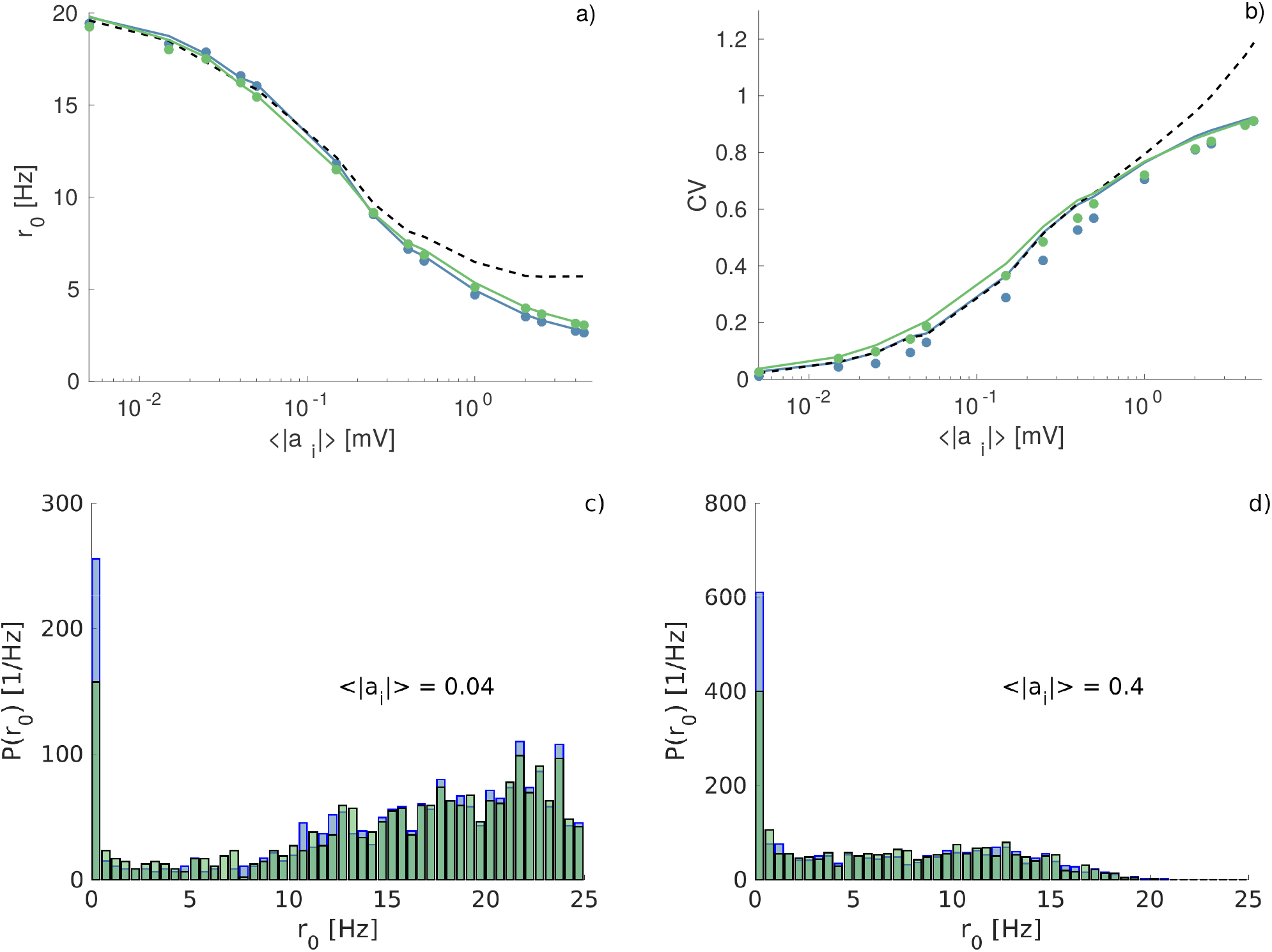
Firing time statistics for heterogenoeus networks as a function of the average synaptic weight. (a) Average network frequency *r̄*_0_ and (b) average coefficient of variation 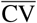 as a function of the average synaptic weight 〈|*a_i_*|〉 for two representative IPSP distributions: DD (blue) and ED (green). Symbols correspond to simulation values and solid lines to the theoretical results from Eqs.(57) and (58). Dashed line denote the results of the DA. The numerically estimated probability distribution functions *P*(*r*_0_) of the single neuron firing rate *r*_0_ are also reported for the two considered IPSP distributions for two values of the average synaptic weight: namely 〈|*a_i_*|〉 = 0.4 (c) and 〈|*a_i_*|〉 = 0.4 (d). Simulations were performed with *N* = 400 neurons, *K* = 20 and a distribution of the input currents uniformly chosen in the interval *μ*(*j*) ∈ [10,11] mV. The time averages are calculated, after discarding an initial transient corresponding to 10^6^ spikes, over the following 10^6^ spikes. Silent neurons (those that do not emit any spike in the considered time lapse) are not included in the statistics. In all cases 〈|*a_i_*|〉 denotes the average value of the corresponding distribution.

As a final remark, we would like to stress that the reported approach works very well also for heterogeneous sparse networks, provided that the collective dynamics is asynchronous. Otherwise, the presence of partial synchronization or of collective oscillations can induce correlations in the input spike trains, which cannot be accounted for with this approach. In particular, to avoid phase locking among the neurons the distribution *Ρ*(*μ*) of the excitabilities should be sufficiently wide.

## Conclusions

In this paper we have reported a theoretical methodology to obtain exact firing statistics for leaky integrate-and-fire neurons subject to discrete inhibitory noise, accounting for Poissonian trains of uncorrelated post-synaptic potentials. Our results represent an extension to generic synaptic weights distribution of the approach developed in^17^ for exponentially distributed post-synaptic potentials. In particular, we report explicit results for the firing rate, the coefficient of variation and the spike train spectrum.

The comparison with numerical simulations reveals a very good agreement for all the considered distributions over all the reported ranges of shot noise amplitude. Moreover, the method is also able to reproduce the average activity of an heterogeneous inhibitory neural network with sparse connectivity, by making use of a self-consistent mean field formulation.

Conversely, the diffusion approximation^3^ (the most used theoretical approach), gives a reasonable estimate of the firing time statistics for sufficiently small IPSP amplitudes, but the agreement rapidly degrades, and reveals large discrepancies already for amplitudes > 0.5 mV, corresponding to physiologically relevant values^8,36,37^.

As a general result we observe that the firing statistics of single neurons is strongly influenced by the shape of the IPSP distributions. Distributions with longer tails lead to smaller firing rate and in general to more regular spike trains (for sufficiently large inhibition). For heterogeneous networks the value of the average IPSP has a noticeable influence on the firing activity of the neuronal population, while the shape of the synaptic weight distributions seem to have a really limited impact on the average properties of the network. However, differences induced by the IPSP distributions are still observable at the level of single neuron.

We have shown that the method works for any choice of IPSP amplitude distributions, however one has to be careful when dealing with the excitatory drive. We have shown that the agreement with numerical simulations is very good when the firing of the neuron is either promoted exclusively by a suprathreshold DC current or provided by the stochastic arrivals of exponentially distributed EPSP amplitudes. In this latter case, an excitatory DC current can as well be present in addition to the EPSP trains, however the DC contribution must be strictly sub-threshold, namely, *μ*_0_ < *v_th_*. Whenever the two firing mechanisms are active at the same time (i.e. *μ*_0_ > *v_th_*), the formulation reported in this paper is no more valid due to the inconsistency in the choice of the proper integration limit *x̄* in Eq.(17). This limitation is illustrated in Fig. 7, where it is reported the response of a LIF neuron, subject to a supra-threshold excitatory DC current *μ*_0_ = 12 mV as well as to Poissonian spike trains of ED excitatory and DD inhibitory synaptic inputs. For small values of the noise intensity, the theoretical approach dramatically fails. In such a region the activity is almost exclusively current driven; i.e, the firing of the neuron promoted by the supra-threshold DC current is much faster than the arrival rate *R_e_* of the EPSPs. This can be confirmed by the fact that in this regime the firing rate coincides with 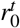 in Eq.(51) for a tonic firing LIF neuron subject to a constant DC current *μ_T_* = *μ*_0_. As the noise intensity grows, corresponding to an increased rate *R_e_* of arrival of the EPSPs, the numerical data approach the theoretical prediction obtained for exponentially distributed EPSPs. The crossover occurs for 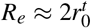, indicating that the firing activity of the neuron is now mainly driven by the stochastic component.

**Figure 7.**
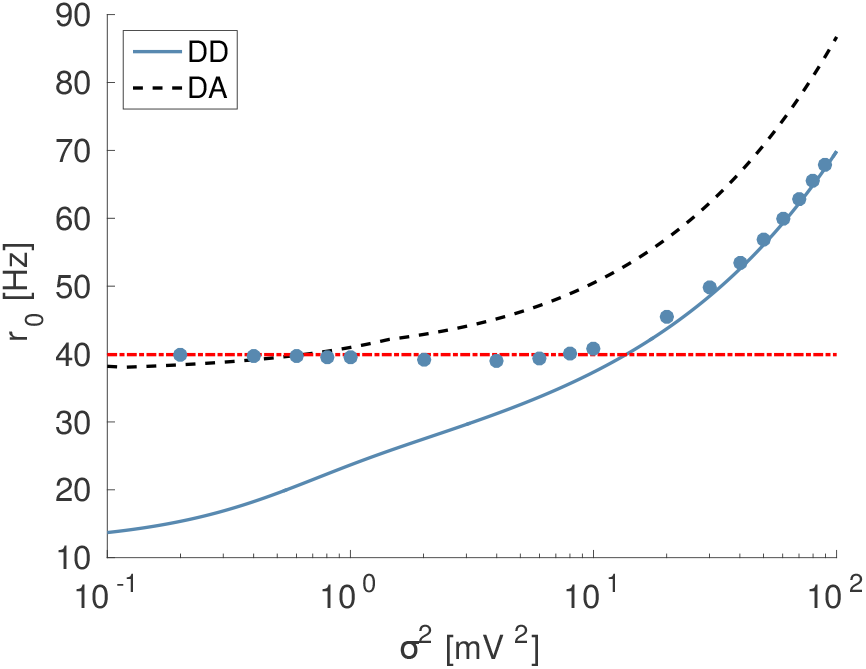
Firing time statistics for exponentially distributed EPSPs and supra-threshold DC current. Firing rate *r*_0_ as a function of noise intensity for excitatory synaptic weights distributed exponentially and a supra-threshold DC current. All parameters as in 5 except for *μ_T_* = *μ*_0_ = 12 mV. The dashed red horizontal line refers to the rate 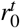 of the LIF neuron subject to constant current *μ_T_* (Eq.(51)). Other symbols and lines have the same meaning as in Fig.2, as well as all the other parameters for the IPSP distribution and the numerical simulations.

**Figure 8.**
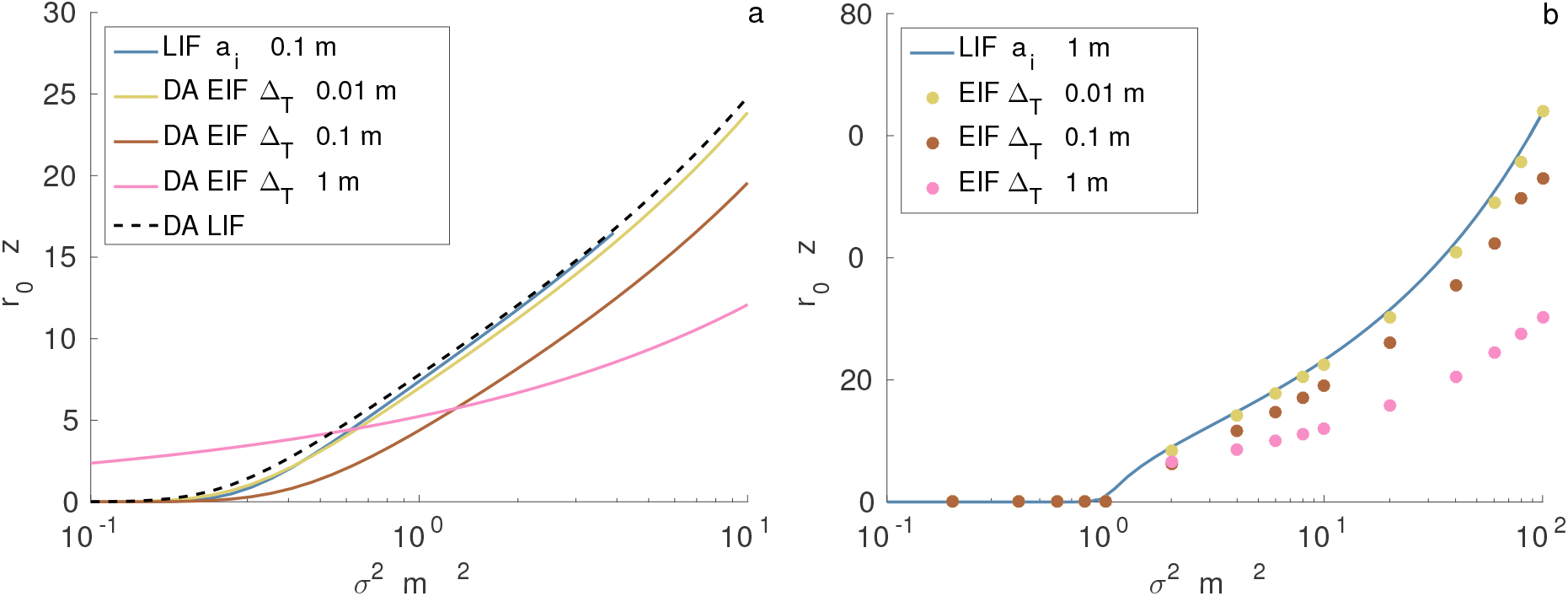
Comparison of the firing rate between LIF and EIF. a) Firing rate in the Diffusion limit for the EIF at different Δ_*T*_ and the comparison with the DA and *δ*-distributed amplitudes of IPSP in the LIF model with small |*a_i_*| = 0.1 mV. Results for the EIF are obtained by solving the stationary state of Eqs.(60) and (61) via the threshold integration method as reported in^3^. For the LIF, the DA and the *δ*-distributed solutions are taken respectively from Eqs.(17) and (51) in the main text. b) Numerical simulations of the EIF in the shot noise case with *δ*-distributed IPSP amplitudes of average |*a_i_*| = 1mV, for the same values of Δ_*T*_ as in panel a), and the corresponding LIF case with the same IPSP distribution. In this panel, the results of the EIF are calculated numerically by integrating Eq. (59) with an Euler scheme with time step *h* = 1 × 10^−3^. When the neuron reaches a large value *v*_∞_ = 80 mV, the remaining time to reach infinity is calculated as *t*_∞_ = *τ* exp((*v_th_* − *ν_∞_*)/Δ_*Τ*_)^1^. In all the cases we have chosen *μ_Τ_* = 9mV, *v_re_* = 5mV, *v_th_* = 10mV and *τ* = 20ms.

An interesting semi-analytic approach has been recently reported for excitatory shot noise in^13^. However, further progresses are required to achieve exact firing time statistics going beyond the diffusion approximation for generic distributions of instantaneous excitatory PSPs, as well as for more realistic neuronal models like the Exponential Integrate and Fire (EIF) neuron^1^ able to reproduce quite accurately the dynamics of cortical neurons^2^. In particular, as shown in the Supplementary Information, the LIF neuron response is a limit case of the EIF dynamics both within the diffusion approximation as well as for shot noise. Furthermore, the diffusion approximation fails also for the EIF to capture the neuronal firing statistics for sufficiently large IPSP amplitudes in particular at low noise intensities.

## Acknowledgments

We thank for useful discussions B. Lindner, G. Mato, A. Politi, G. Martelloni, MJE Richardson, M. Timme. This work has been partially supported by the European Commission under the program “Marie Curie Network for Initial Training”, through the project N. 289146, “Neural Engineering Transformative Technologies (NETT)” and by the A*MIDEX grant (No. ANR-11- IDEX-0001-02) funded by the French Government “Programme Investissements d’Avenir” (D.A.-G. and A.T.). AT gratefully acknowledges a “VELUX Visiting Professor Grant” from Villum Fonden that supported his stay at Aarhus University during the initial stages of this work in 2011. AI is supported by the Danish Council for Independent Research and the Villum Fonden. This work has been completed at Max Planck Institute for the Physics of Complex Systems in Dresden (Germany) as part of the activity of the Advanced Study Group 2016 entitled ”From Microscopic to Collective Dynamics in Neural Circuits”.

## Author contributions statement

S.O and D.A-G performed the numerical simulations. D.A-G and A.T prepared the manuscript. All the authors developed the theoretical methods and reviewed the manuscript.

## Additional information

**Competing financial interests** Authors declare to have no competing financial interests that might have influenced this manuscript.

## Supplementary Information

### Exponential Integrate and Fire

The Exponential Integrate-and-Fire (EIF) model is a simple non-linear integrate and fire neuronal model introduced by Fourcaud-Trocmé et al.^1^ able to reproduce quite accurately the dynamics of cortical neurons^2^. The model can be written as follows

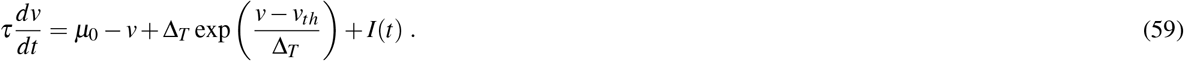

where *μ*_0_ represents an external DC current and *I*(*t*) the synaptic drive. In this model, the spike generation occurs in a finite time controlled by the parameter Δ_*T*_. In particular, once the membrane potential has reached the threshold value *v_th_* this will rapidly grow towards infinity in a finite time interval, the parameter Δ_*Τ*_ establishes how fast the infinite limit is reached. In the limit *Δ_T_* → 0, the spike generation is instantaneous and the LIF model is recovered. As in the usual LIF model, once the neuron has fired its membrane potential is resetted to the value *v_re_* = 5 mV, we also set *τ* = 20 ms and *v_th_* = 10 mV, as in the LIF model studied in the article.

As a first analysis, we will examine the response of the EIF neuron subject to small Gaussian noise of zero average and intensity *σ*, this can be obtained by solving the associated continuity equation for the probability *P*(*v*, *t*) of finding the membrane voltage between *v* and *v* + *dv* at time *t*, which reads as^1^:

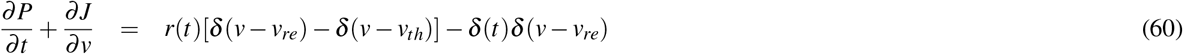

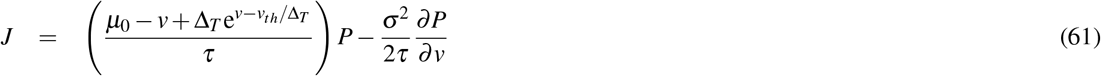
where *J = J*(*v*, *t*) is the associated flux.. In this case, since the effective threshold is located at infinity, the steady firing rate can be evaluated as

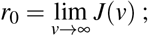
where *J*(*v*) is the stationary solution of the continuity equation for the flux.

In particular, we made use of the threshold-integration method^3^ to calculate the firing rate *r*_0_ of the EIF neuron subject to inhibitory inputs and compare it with the diffusion approximation (DA) for the LIF and the corresponding shot noise solution for *δ*-distributed IPSP amplitudes with |*a_i_*| = 0.1 mV. The results are shown in 8 (a), where it is clearly shown that for Δ*_Τ_* → 0 the EIF results converge to the LIF solution, both for the DA and the the shot noise results found for small IPSP. Furthermore direct simulations of the EIF and the corresponding shot noise solution of the LIF for large IPSP amplitudes, namely |*a_i_*| = 1 mV, show that also in this case the LIF limit is recovered for Δ*_T_* → 0. However it is clear that also in the case of the EIF, the diffusion limit is unable to capture the onset of the activity and the firing rate at small intensities.

